# Genome-wide profiling of an enhancer-associated histone modification reveals the influence of asthma on the epigenome of the airway epithelium

**DOI:** 10.1101/282889

**Authors:** Peter McErlean, Audrey Kelly, Jaideep Dhariwal, Max Kirtland, Julie Watson, Ismael Ranz, Alka Saxena, David J. Cousins, Roberto Solari, Michael R. Edwards, Sebastian L. Johnston, Paul Lavender, on behalf of the MRC-GSK Strategic Alliance Consortium

**Affiliations:** Peter Gorer Department of Immunobiology, King’s College London, London, UK; MRC Asthma UK Centre in Allergic Mechanisms of Asthma, London, UK; Airway Disease Infection Section, National Heart and Lung Institute, Imperial College London, UK; Biomedical Research Centre at Guy’s and St Thomas’ NHS Foundation Trust.; NIHR Respiratory Biomedical Research Unit, Department of Infection, Immunity & Inflammation, Leicester Institute for Lung Health, University of Leicester, Leicester, UK; Airway Disease Infection Section, National Heart and Lung Institute, Imperial College London, London, UK; MRC and Asthma UK Centre in Allergic Mechanism of Asthma, London, UK; GlaxoSmithKline, Allergic Inflammation Discovery Performance Unit, Respiratory Therapy Area, Stevenage, UK; Department of Respiratory Medicine & Allergy, King’s College London, London, UK; Guy’s and St Thomas’ NHS Trust, London, UK

**Keywords:** Asthma, Epigenomics, Super Enhancers, Histone, Chromatin, ChIP-Seq, Epithelium, Airways, Respiratory

## Abstract

**Background:** Asthma is a chronic airway disease driven by complex genetic-environmental interactions. The role of epigenetic modifications in bronchial epithelial cells (BECs) in asthma is poorly understood. We undertook genome-wide profiling of the enhancer-associated histone modification H3K27ac in BECs from people with asthma (n=4) and healthy controls (n=3).

**Results:** We identified n=4,321 (FDR <0.05) regions exhibiting differential H3K27ac enrichment between asthma and health clustering at genes associated predominately with epithelial processes (EMT). Asthma dramatically influenced the BEC enhancer landscape and we identified asthma-associated Super-Enhancers encompassing genes encoding transcription factors (*TP63*) and enzymes regulating lipid metabolism (*PTGS1*). We integrated published datasets to identify epithelium-specific transcription factors associated with H3K27ac in asthma (*TP73*) and identify initial relationships between asthma-associated changes in H3K27ac, DNA methylation, genetic susceptibility and transcriptional profiles. Finally, we used a CRISPR-based approach to functionally evaluate components of the H3K27ac-asthma landscape *in vitro* and provide proof of principal that asthma-associated gene expression (*TLR3*) is driven in part by aberrant histone acetylation.

**Conclusion:** Our small study validates the combination of genome-wide and epigenome-editing approaches in deciphering the molecular mechanisms underlying asthma pathogenesis.

## Background

Asthma is a chronic inflammatory disease of the airways affecting over 230 million people worldwide(1). Driven by complex genetic-environmental interactions, asthma’s origins, triggers and clinical presentations are heterogeneous, posing challenges to understanding disease development, molecular components and the generation of more effective therapies(2).

The airway epithelium provides an initial physical barrier and biological responses to environmental insults. The airway epithelium of people with asthma is characterized by altered phenotypic and transcriptional characteristics including excessive mucus production, defects in antiviral responses and repair(3–5) and the predominance of signals associated with type 2 (T2) inflammation(6, 7). However, little is known about the molecular mechanisms that underpin the transcriptional profiles of airway epithelium in asthma.

The accessibility of areas within DNA contributing to transcription (transcription factor binding sites, regulatory elements/enhancers) is determined in part by epigenomic mechanisms including DNA methylation and histone modifications. Chromatin-immunoprecipitation coupled with high-through-put sequencing (ChIP-Seq) has revealed specific modifications on histone H3 are associated with various transcriptional states and genomic features. For example while occurring at transcriptional start sites (TSS’s), H3 Lys4 dimethylation (H3K4me2) and H3 Lys27 acetylation (H3K27ac) are enriched predominately in intergenic (i.e. non-coding) regions and demarcate poised and active enhancers respectively(8).

Studies of epigenomic mechanisms have determined the epigenomes across a variety of cell types is altered in asthma(9–11). ChIP-Seq profiling of H3K4me2 in CD4^+^ T cells from individuals with asthma has identified putative asthma-associated enhancers(10), supporting the use of genome-wide approaches to identify novel epigenomic mechanisms in asthma. However, while DNA methylation has been broadly investigated(12–14), genome-wide investigations of histone modifications in the airway epithelium in asthma have yet to be undertaken.

Ranking ChIP-Seq signals revealed that regions of the genome exhibit increased enrichment of enhancer-associated histone modifications in a cell-type and disease-specific manner. These regions, termed ‘Super-Enhancers’ (SEs) differ from otherwise ‘typical’ enhancers by exhibiting sustained enrichment across several kilobases (kb), encompass ‘master’ transcription factors (TFs) important in cell-identity and are more likely to harbor disease-associated single nucleotide polymorphisms (SNPs)(15, 16). Super-Enhancers have been identified across numerous cell-types and diseases and are of potential interest for therapeutic intervention(17–19). However, airway epithelial cell and asthma-associated SEs have yet to be described.

Determining the functional consequences of epigenomic changes has been bolstered by the recent developments of the clustered, regularly interspaced, short palindromic repeats (CRISPR)-Cas9 system(20). By fusing catalytic domains of epigenome-modifying factors (e.g. DNA methyltransferases(21) and histone acetyltransferases(22)) to the catalytically-dead Cas9 (dCas9), site-specific epigenome editing can be achieved in a guide RNA (gRNA)-mediated manner and the consequence of epigenome alterations on gene transcription determined. Using dCas9-based editing, it is now possible to recapitulate components of disease-associated epigenomes *in vitro*, providing a means to accurately determine the contribution of aberrant epigenomic landscapes to disease-associated transcriptional profiles.

In the current study, we profiled H3K27ac via ChIP-Seq in bronchial epithelial cells (BECs) from people with asthma and healthy individuals. We undertook genome-wide analysis and identified regions of differential H3K27ac in asthma and by profiling an enhancer-associated histone modification, were able to identify airway epithelial and asthma-associated SEs. We additionally identified epithelial-specific TFs associated with the altered H3K27ac landscape in asthma and relationships between asthma-associated H3K27ac and DNA methylation, genetic susceptibility and transcriptional profiles. Finally, we demonstrate by recapitulating the epigenomic landscape via a dCas9-based approach, that aberrant histone acetylation in asthma can drive asthma-associated gene expression.

## Results

### Asthma influences H3K27ac enrichment in airway epithelial cells

To investigate the influence of asthma on the epigenome of the airway epithelium, we undertook genome-wide profiling of H3K27ac via ChIP-Seq in BECs from healthy controls (n=3) and adults with asthma (n=4, Table S1). We identified regions exhibiting the greatest H3K27ac enrichment for each volunteer (peaks, Table S2) and then by comparing enrichment across a consensus peak set identified n=4,321 differentially enriched regions (DERs, FDR<0.05, DiffBind) between asthma and healthy BECs (Table S3).

Initial PCA confirmed that the greatest variation in DERs was observed between study groups, indicating that despite *ex vivo* culture, differences in the epigenome were present in BECs (Figure 1A). Asthma DERs encompassed regions with a Gain (n=3,061) or Loss (n=1,260) in H3K27ac enrichment (Figure 1B) and occurred predominately at distal intergenic and promoter/intronic sites respectively (Figure 1C).

**Figure 1:**
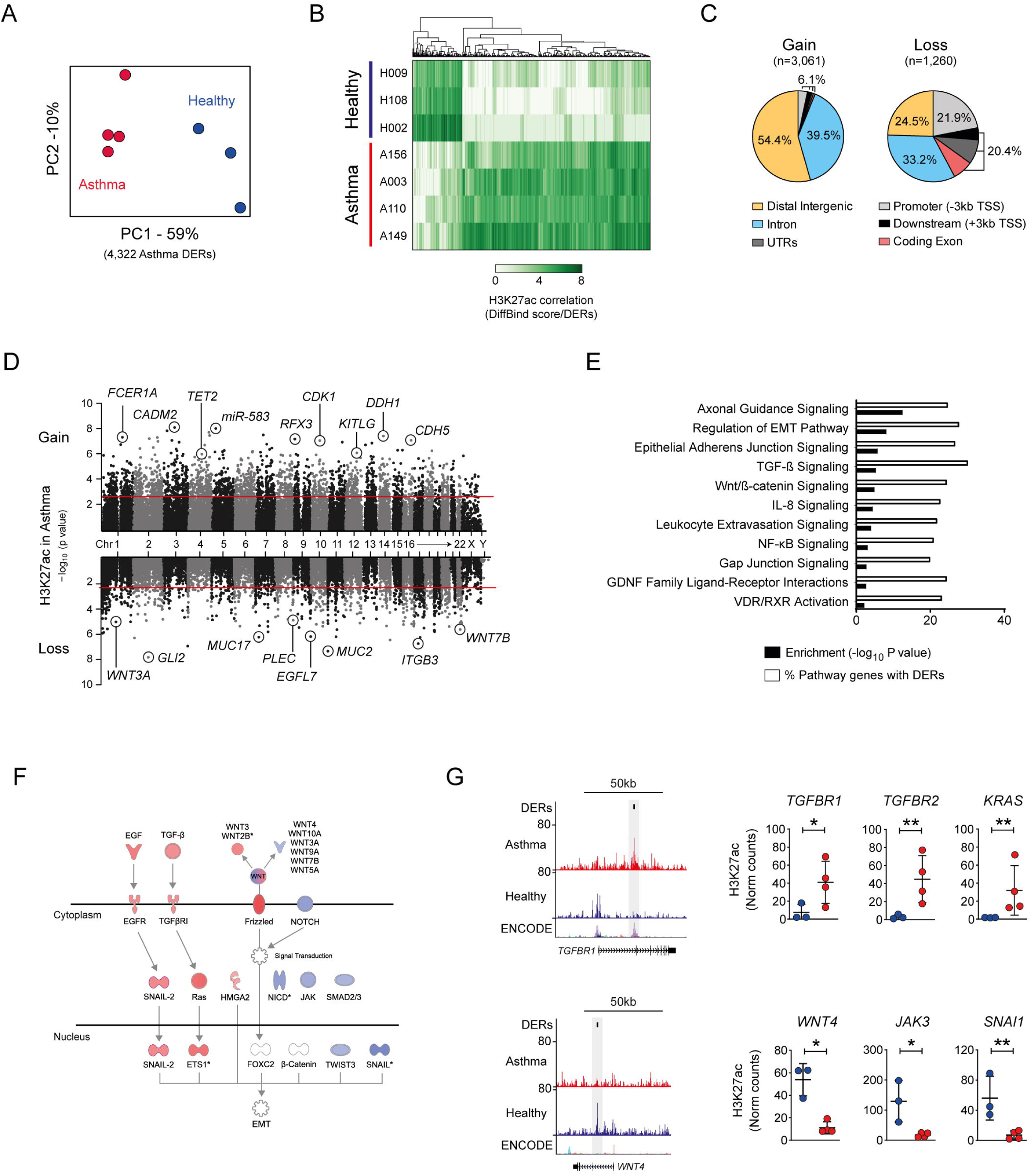
Influence of asthma on the epigenome of the airway epithelium. (A) Principal component analysis of H3K27ac across differentially enriched regions (DERs) identified by DiffBind. Clustering identifies variation both between and within healthy (blue) and asthma (red) BECs. (B) Heatmap depicting volunteer correlation across asthma DERs and the presence of gain and Loss of H3K27ac in asthma BECs. (C) Distribution of Asthma DERs across genomic features. As H3K27ac is associated with enhancers, most DERs occurred in distal/intergenic sites. TSS – transcriptional start site, UTR – un-translated region. (D) Manhattan plot depicting genome-wide distribution of DERs. Each dot represents an Asthma DER (i.e. 500bp window) plotted in relation to its location and statistical significance. Dashed line represents FDR *P*<0.05. (E) DER-associated genes are enriched in pathways associated with epithelial biology and asthma pathophysiology (-log_10_ *P* values, black bars). Pathway analysis also revealed up to 30% of genes within the same pathway exhibited differential H3K27ac in asthma (% of pathway genes with DERs, white bars). (F). Condensed versions of the epithelial-to-mesenchymal transition (EMT) pathway depicting how numerous components within the same pathway gain (red) and lose (blue) H3K27ac in asthma BECs. (G) Genome tracks depicting H3K27ac enrichment in BECs across example Gain (*TGFBR1*) and Loss asthma DERs (*WNT4*). Data tracks from bottom to top; genes (RefSeq annotation), layered H3K27ac from ENCODE, merged data from healthy (n=3, blue) and asthma BECs (n=4, red) and Asthma DERs (black boxes). Dot plots depict enrichment across volunteers at select genes with gain and loss DERs (\**P*<0.05, \*\**P*<0.01,DiffBind). Asthma=red, healthy=blue.

Because our asthmatic volunteers were on inhaled corticosteroids (ICS), we investigated if asthma DERs could be the result of treatment-induced glucocorticoid receptor (GR) binding. We observed 2.3% of asthma DERs (n=101, P=0.076 Fishers exact test) overlapped with GR-binding sites identified in dexamethasone-(DEX)-treated airway epithelial cells(38), suggesting that glucocorticoid treatment was not a primary cause of epigenome reorganization in these subjects (Figure S1A-B).

Although occurring at intergenic sites genome-wide, asthma DERs were proximal to n=3,062 genes which predominately encoded enzymes and transcriptional regulators (Figures 1D and S1C). Pathway analysis revealed DER-associated genes contributed to epithelial (e.g. cellular adherens) and asthma-associated processes (e.g. TGF-β signaling) with a up to 30% of all genes within these pathways exhibiting differential H3K27ac in asthma (range 19.9%-30.1%, Figure 1F-G). Taken together, these data reveal that the airway epithelium in asthma is characterized by dynamic and distinct changes in the histone landscape.

### Epithelial and asthma-associated Super-Enhancers are associated with enzymes involved in lipid metabolism

Since we profiled an enhancer-associated histone modification, we sought to determine the enhancer landscapes the airway epithelium (H3K27ac enrichment >2kb from TSS’s). H3K27ac peaks identified in merged datasets from healthy and asthma BECs were ranked by ChIP-Seq signal (ROSE) to determine typical- and super-enhancer landscapes (SEs, Figure 2A). Given SEs can characterize cell-types and disease, we focused on the SEs specifically and observed several shared and unique SEs in this initial analysis. We subsequently identified and compared SEs from each volunteer revealing the presence of Common (airway cell-specific), Healthy and Asthma-associated SEs in BECs (Figures 2B, Table S4).

**Figure 2:**
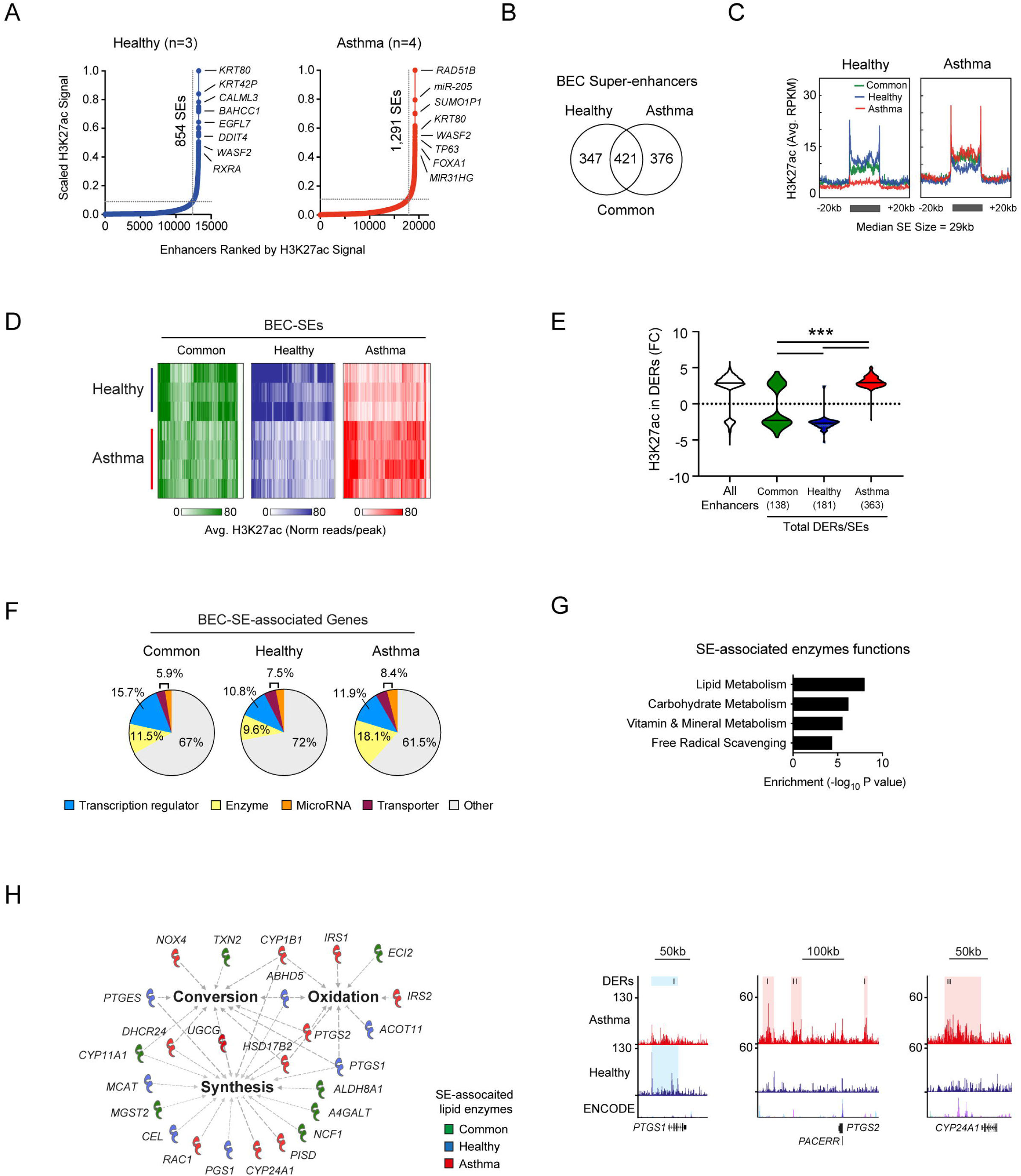
Identification of cell-specific and asthma-associated super enhancers in the airway epithelium. (A) Ranking of H3K27ac ChIP-Seq signal from merged Healthy (blue, n=3) and asthma BECs (red, n=4) identified super enhancers in BECs (BEC-SEs) and associated genes. BEC-SEs were shared (e.g. *KRT80*) and unique to healthy (e.g. *CALML3*) or asthma (e.g. *RAD51B*). (B) Comparative analysis of SEs from individual volunteers identified three categories of BEC-SEs; those common across all volunteers (airway cell-specific) and those associated with healthy or asthma BECs (see methods and Table S4). (C) ‘Meta gene’ plots summarizing H3K27ac enrichment (RPKM) in healthy and asthma BECs across Common, Healthy- and Asthma-associated SEs. (D) Heatmap depicting H3K27ac enrichment for each volunteer across BEC-SE categories. (E) Plot depicting log fold change (FC) of all asthma DERs located within BEC-SEs. Asthma BECs exhibited marked changes in H3K27ac across airway cell-specific (i.e. common, green) and healthy- (blue) or asthma- (red) SEs (median ±max-min values, \*\*\**P*<0.001, ANOVA/ Tukeys multiple comparisons). (F) Consistent with super enhancers in other cells types, BEC-SE-associated genes included transcriptional regulators (blue). However, we also identified substantial numbers of BEC-SE-associated genes encoded enzymes (yellow). (G) Pathway analysis indicating that BEC-SE-associated enzymes were enriched in various metabolic processes, particularly those associated with lipids. (H) Lipid-associated processes (bold) and enzymes encompassed by BEC-SEs. Genome tracks additionally depict H3K27ac enrichment, DERs and location of BEC-SEs across select lipid-associated enzymes.

Like other SEs, BEC-SEs spanned large genomic regions with increased enrichment (Figure 2C) and were most closely related to SEs identified in mucosal cell-types and tissues (e.g. oesophagus, mammary epithelium, Figure S1D). However, similarity did not exceed 54.3% suggesting the BEC-SEs we identified were airway-cell specific. Similarly, study volunteers exhibited H3K27ac enrichment consistent with BEC-SE category, suggesting our BEC-SEs were disease-specific (Figure 2D).

We next investigated differential H3K27ac in BEC-SEs found 39.3% of BEC-SEs contained DERs (range 1-6 DERs/SEs) covering on average only 7% of the total SEs size (range 0.8%-79.5%; Table S4). However, while jointly typical- and super-enhancers overlapped 79.2% of asthma DERs (Figure S1E), plotting fold change of DERs across all enhancers identified dramatic changes in H3K27ac in SEs. Specifically, Asthma BECs exhibited marked decreases in H3K27ac in Common- and Healthy-SEs and increased H3K27ac in Asthma-SEs (Figures 2E).

Consistent with SEs encompassing loci expressing ‘master’ TFs, we found between 10.8-15.7% of BEC-SE-associated genes encoded transcription regulators (Figure 2F), more so than typical enhancers (8.2%, Figure S1F). However, we additionally found BEC-SE-associated genes encoded enzymes (Table S4). Indeed, a greater proportion of asthma-SE-associated genes encoded enzymes than transcriptional regulators (18.15% vs. 11.9% respectively, Figure 2F). BEC-SE-associated enzymes were enriched in metabolic processes, particularly the conversion, synthesis and oxidation of lipids (Figure 2G). Further investigation revealed several lipid-associated enzymes implicated in asthma pathogenesis to be encompassed by asthma-SEs (e.g. *PTGS1, PTGS2, PTGS1*, Figure 2H). Taken together, these data indicate that asthma has a dramatic influence on the location and enrichment of SEs in the respiratory epithelium.

### Transcription factors associated with H3K27ac in asthma

We next expanded on transcription regulators and identified enrichment of TF binding motifs in Asthma DERs and BEC-SEs. Enriched motifs were predominately from members of the p53-related, forkhead-box and AP1-families, TFs selectively enriched in individual cell types of bronchial epithelium (e.g. P63) and aberrantly expressed in inflamed tissues respectively (e.g. Jun/Fos, Figure 3A). We refined computational predictions (see methods) and found despite association with epithelial process (n=201, Figures S2A-B), most TFs identified had protein expression across many tissue types. However, our approach identified a subset of TFs with protein expression predominately in the respiratory epithelium (e.g. TP63, Figure 3B).

**Figure 3:**
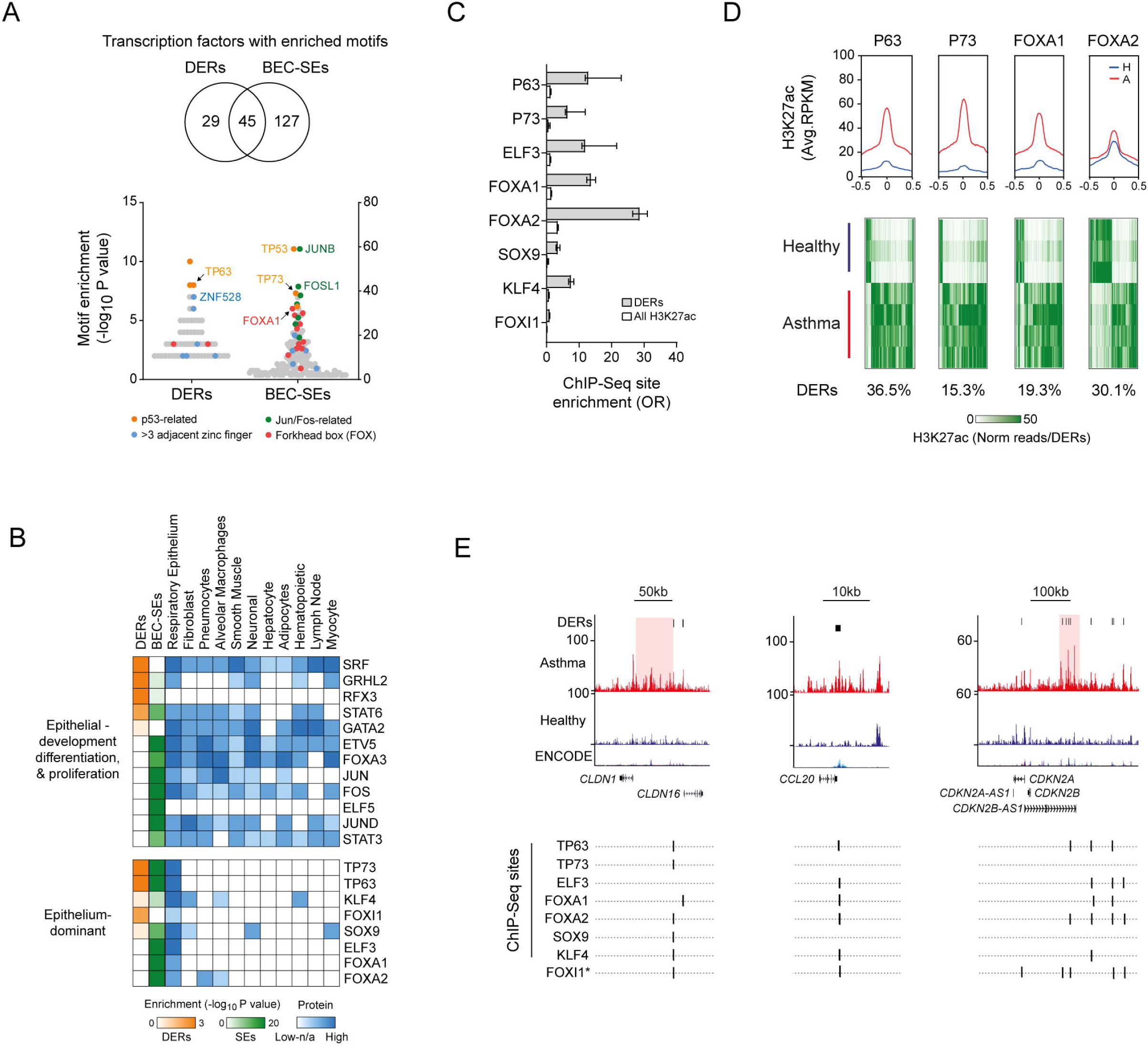
Identification of airway epithelium-dominant transcription factors (TFs) associated with H3K27ac in asthma. (A) Enrichment of TF motifs across asthma DERs and BEC-SEs. Examples of the most enriched and prevalent TF families are indicated. (B) Heat map depicting motif enrichment, protein expression and tissue distribution of TFs enriched in asthma DERs and BEC-SEs. Most enriched TFs were expressed across many tissue types. However, a subset were expressed almost exclusively in the respiratory epithelium (bottom panel). (C) Enrichment of DERs at bona-fide epithelium-dominant TFs binding sites identified via ChIP-Seq. All enrichment in DERs and H3K27ac in BECs were significant (P<0.0001, Fishers odds ratio +/-95% CI). (D) H3K27ac profiles of healthy (blue) and asthma (red) BECs at epithelium-dominant TF binding sites identified via ChIP-Seq. Plots represent enrichment ±0.5kb around binding site center. Heatmaps depict H3K27ac enrichment for each DERs containing epithelium-dominant TF binding sites across study volunteers. Proportion of total DERs with sites indicated below (see also Figure S2E) (E) H3K27ac enrichment and ChIP-Seq-validated or predicted (*) epithelium-dominant TFs binding sites across epithelial, cell recruitment and cell cycle-associated genes. *CLDN1* and *CDKN2B-AS1* are encompassed by asthma-associated SE’s (red).

We focused on Epithelium-dominant TFs and found those encompassed by BEC-SEs (Figure S2C) have been previously implicated in mouse models of allergic inflammation (*FOXA2*)(39, 40), asthma, nasal polyposis and mucociliary development (*TP63, TP73*)(41–44). We sought to validate computational predications further (Figure S2D) and found DERs were enriched at bona-fide epithelial-TF sites as determined by ChIP-Seq (Figure 3C). Profiling revealed Asthma BECs on average exhibited more H3K27ac at epithelial-TF ChIP-Seq sites, encompassing up to 36.5% of asthma DERs (Figure 3D and S2E). However, plotting H3K27ac enrichment across study volunteers indicated both gain and loss at Epithelium-dominant TF sites.

Finally, we found Epithelium-dominant TF binding sites co-localized at DERs populating genes associated with epithelial integrity (e.g. *CLDN16*), inflammatory cell recruitment (e.g. *CCL20*) and cell cycle (e.g. CDKN2A, Figure 3E). These findings suggest that asthma-associated changes in the histone landscape are accompanied by binding of airway cell-specific and otherwise ubiquitously expressed TFs.

### Asthma-associated changes in H3K27ac and DNA methylation

Since H3K27ac represents only one of many epigenomic mechanisms, we investigated how our findings related to other epigenetic features readily studied in asthma. We focused on the relationship between asthma DERs and DNA methylation and found dramatic differences CpG content in DERs and H3K27ac enrichment in relation to CpG islands (Figure S1G-H), likely reflecting Loss DERs (21.9%) and CGI’s co-occurring at promoter regions (see Figure 1C).

Considering these differences, we investigated the relationship between asthma DERs, BEC-SEs and asthma-associated changes CpG methylation identified in the airway epithelium(13). We found 6.8% of asthma DERs (n=295) and 53.5% BEC-SEs (n=613) were enriched for differentially methylated CpGs (Figure 3A, P<0.001, Fishers exact Test Vs all diff meth CpGs) and a weak but significant correlation between asthma-associated changes in H3K27ac and CpG methylation (Figure 3B). Interestingly, while <1% (n=358) of asthma-associated CpG methylation overlapped asthma DERs, 4.0% (n=1,653) resided within Common- and Healthy-SEs predominately at genes involved in epithelial processes (e.g. *NOTCH1*, *PLEC*, Figure 4C). Taken together, these data suggest concurrent changes in epigenomic mechanisms in asthma BECs.

**Figure 4:**
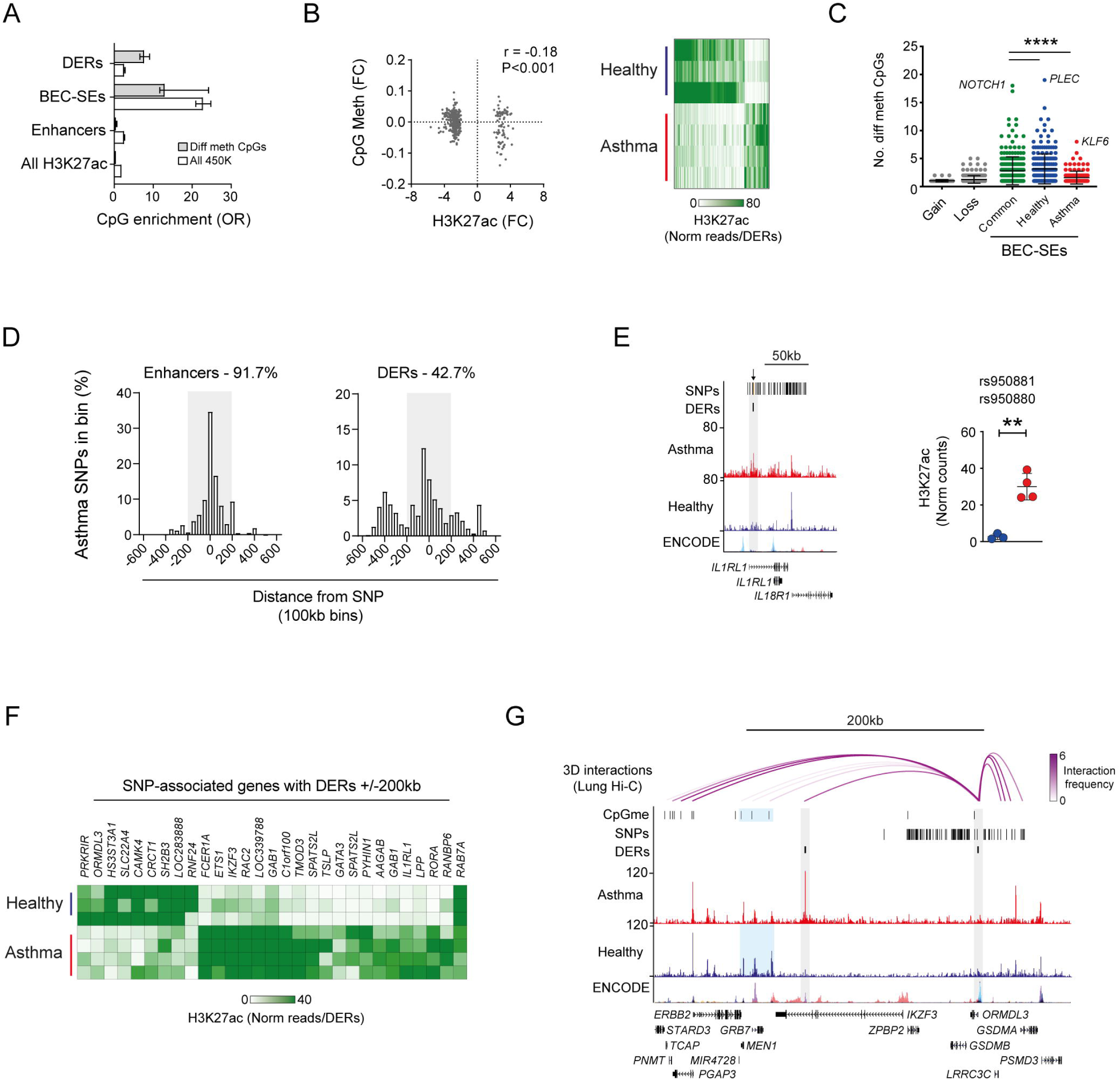
Relationship between H3K27ac, DNA methylation and asthma SNPs in the airway epithelium. (A) Enrichment of differentially methylated CpGs identified by Nicodemus-Johnson *et. al*(13) across asthma DERs and other features of H3K27ac in BECs (all enrichment were significant P<0.0001, Fishers odds ratio +/-95% CI). (B) Correlation between changes in CpG methylation and H3K27ac in asthma BECs. Heatmap depicts H3K27ac enrichment for DERS overlapping differentially methylated CpGs across all study volunteers (FC – log fold change). (C) Asthma-associated changes in CpG methylation were more likely to occur in Common and Healthy SEs. BEC-SE-associated genes with the greatest density of differentially methylated CpGs are indicated (**** *P*<0.0001, ANOVA). (D) Graph summarizing overlap and distance between asthma SNPs, BEC enhancers and DERs. Percentage of total SNPs within ±200kb (greay box) of each feature are indicated. (E) Genome tracks and dot plot depicting H3K27ac of DERs overlapping asthma SNPs at *IL1RL1* (orange lines and arrows, top track) in healthy (blue) and asthma (red) BECs. (\*\**P*<0.01, DiffBind). (F) Heatmap depicting gain and loss of H3K27ac enrichment for DERs within 200kb of an asthma-associated SNP. For clarity, SNPs are identified by their associated gene. (G) Genome tracks depicting H3K27ac in BECs and long-range chromosomal interactions in lung tissue (top tracks) across the 17q21 locus. Purple arcs depict interactions that join the *ORMDL3* TSS with asthma SNPs, DERs and differentially methylated CpGs (CpGme) in three-dimensional space as determined by proximity assays (Hi-C, 5kb bins). Interactions are colored according to distance-normalized interaction frequency. NB: *ORMDL3* is encoded on the negative strand, so features to the right and left are up- and downstream of the TSS respectively.

### Interactions between Asthma-associated H3K27ac and SNPs are mediated via 3D chromatin architecture

Since previous studies have identified enrichment of disease-associated SNPs in enhancers(10), we looked at the relationship between asthma SNPs and H3K27ac in BECs. Asthma-associated SNPs overlapped and were enriched in BEC enhancers (n=23 SNPs/28 enhancers, Figure 4D and S1I). However, direct SNP overlap with DERs and BEC-SEs occurred only at *IL1RL1* (encoding the IL33 receptor ST2, Figure 4E)(45) and with *IL6R* (Common-SE) and *C1orf100* (Asthma-SE) respectively (data not shown). As a result, we did not observe a statistical enrichment of asthma SNPs in DERs or BEC-SEs (Figure S1I). We found that while most asthma SNPs were found within 200kb of a BEC enhancer, only 42.7% were within the same distance of an asthma DER (Figure 4D). However, we observed highly consistent changes in H3K27ac across study volunteers for those DERs within 200kb of an asthma SNP (Figure 4F).

Since most SNPs reside within non-coding regions and influence gene expression through long-range chromosomal interactions (13, 46), we investigated if SNPs and DERs are linked by 3D chromatin architecture. We focused on a DER proximal to *ORMDL3* TSS, a key site of the asthma susceptibility locus 17q21 and found that long-range interactions connect the *ORMDL3* DER to asthma SNPs upstream at *GSDMA* (Figure 5G). Furthermore, interactions could link DERs, BEC-SEs and CpGs with differential methylation in asthma downstream at *IKZF3, GRB7/MEN1* and *ERBB2* respectively. Taken together, these data suggest that 3D chromatin interactions links disease-associated polymorphisms and epigenomic changes in asthma.

**Figure 5:**
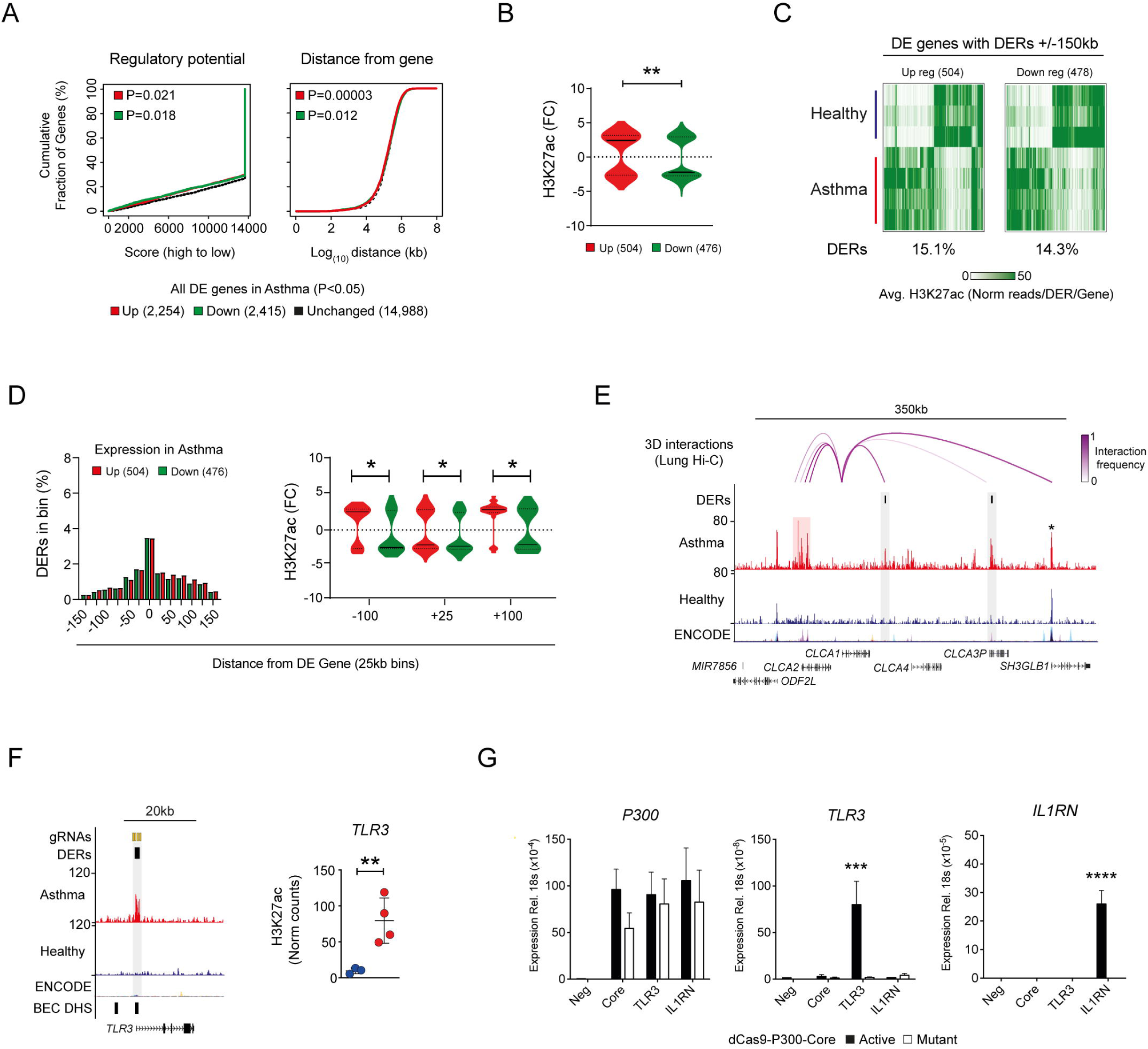
Relationship between H3K27ac and gene expression in asthma and recapitulation of the H3K27ac landscape *in vitro*. (A) Plots generated by BETA analysis depicting the relationship between DERs and differential gene expression (DE) in asthma. Overall, gain and loss of H3K27ac was associated with transcriptional activation (up regulated genes - red) and repression (down regulated genes - green) in asthma respectively. However, this was more strongly related to distance of DERs to DE genes. (B) Plots showing distribution of H3K27ac enrichment for all DERs associated with DE genes identified in BECs. (\*\**P*<0.01, Kolmogorov-Smirnov test). (C) Heatmap depicting both gain and loss of H3K27ac across study volunteers for DERs within 150kb of DE genes up and down regulated in asthma. (D) Graph summarizing number of asthma DERs associated with DE genes by distance (25kb bins) and direction of gene expression in asthma (i.e. up or down regulated). Plots showing changes in H3K27ac enrichment are evident at selected regions up and downstream from DE gene TSSs (\**P*<0.05, Kolmogorov-Smirnov test). (E) Genome tracks depicting H3K27ac in BECs and long-range chromosomal interactions in whole lung (top track) across the *CLCA1* locus. Although exhibiting minimal H3K27ac in BECs, *CLCA1* interacts with asthma DERs spanning ∼200kb (purple arcs) including those near *CLCA3P* and an asthma-SE at *CLCA2* (highlighted in red). Note no change in H3K27ac is evident at *SH3GLB1* (star). (F) Genome tracks depicting H3K27ac across *TLR3* and location of DER-specific gRNAs (top track - orange lines). Regions of ‘open chromatin’ identified in BECs are depicted in the bottom track (DNase hypersensitivity sites - DHS). Dot plots depicting H3K27ac enrichment of DER targeted by gRNAs across healthy (blue) and asthma volunteers (\*\**P*<0.01, DiffBind). (G) Using a CRISPR-based approach, acetylation (dCas9-P300-Core) was targeted to an asthma-DER at *TLR3* using pooled guide RNAs. Increased P300 indicated successful delivery and expression of dCas9 constructs across all gRNA pools. However, *TLR3* gene expression was selectively induced in a DER-specific gRNA and acetylation-dependent manner. A pool of non-DER-targeting gRNAs to *IL1RN* were used as control. Core - only P300 constructs transfected (n=3, Mean ±SEM, *** *P*<0.001, ANOVA/Tukeys multiple comparisons).

### Alterations to H3K27ac are related to asthma-associated transcriptional profiles

Since histone acetylation is associated with transcriptional activation, we investigated the relationship between H3K27ac and gene expression in asthma (34). Using BETA analysis (see methods), we determined that changes in H3K27ac had an overall small association with asthma-associated gene expression. However, this relationship was more pronounced when distance of DERs from differentially expressed (DE) genes was considered (Figure 5A).

As evidenced by BETA analysis, there was a difference in enrichment of H3K27ac at DE genes with up and down regulated genes containing more gain and loss DERs respectively (Figure 5B). However, when looking at enrichment across study volunteers we observed discordance in the relationship between H3K27ac and gene expression with the presence of both gain and loss DERs at DE genes (Figure 5C).

To clarify this discordance, we looked closer at the relationship between distance of DERs to DE genes and expression in asthma. While there was no clear difference in total number of DERs by distance, enrichment of H3K27ac at DE genes were confined to regions up and down stream of gene TSSs (e.g. Up regulated genes gained more H3K27ac 100kb downstream of their TSS, Figure 5D).

To further elucidate the relationship between genomic distance, DERs and gene expression (Figure 1C), we investigated if like SNPs; DERs are connected to DE genes via long-range interactions. We focused on the locus of the Th2-high signature gene *CLCA1*(47) and found that while there was minimal H3K27ac enrichment in either healthy or asthma BECs across *CLCA1* itself, we observed interactions between *CLCA1* and asthma DERs located downstream (Figure 5E). An additional interaction was also evident with an asthma-SE located upstream and encompassing *CLCA2*. Taken together, these data reveal complexity between the histone landscape and gene transcription and further implicates 3D architecture in asthma-associated gene expression.

### Transcriptional consequences of deposition of H3K27ac in asthma

Finally, we employed a CRISPR-based epigenome editing approach to recapitulate the epigenome observed in asthma BECs and further clarify the relationship between H3K27ac and gene expression in asthma. We focused on *TLR3*, a key gene in the antiviral response whose transcriptional start site was largely devoid of H3K27Ac in healthy subjects yet exhibited a gain in H3K27ac in asthma BECs (Figure 5F)). We investigated the effects of acetylation on *TLR3* gene transcription in human embryonic kidney cells (HEK293T), this cell line has little or no endogenous *TLR3* expression and as such mirror’s expression of the gene in healthy BECs.

Using dCas9 fused with the catalytic core of histone acetyltransferase P300 (dCas9-P300-Core)(22), acetylation was targeted via gRNAs to DERs gaining H3K27ac at the *TLR3* TSS (Figure 5F). We found *TLR3* gene expression could be selectively induced using DER-specific gRNAs only in the presence of the active form of dCas9-P300-Core (Figures 5G). Additionally, we found these acetylation-dependent transcriptional effects could be mediated using either pooled and single DER-specific gRNAs (Figure S2F). These data indicate that deposition of a single epigenetic mark, namely differential H3K27ac in asthma, can drive gene expression from a non-expressed locus.

## Discussion

Our study sought to establish whether we could observe differences in epigenetic landscapes in BECs from asthmatic volunteers compared to healthy counterparts. Since it is the best indicator of transcriptional regulatory domains and enriches at genomic features such as TSSs and intergenic regions/enhancers, we studied the effects of asthma on H3K27ac distribution across the genome and identified differential H3K27ac enrichment in asthma BECs, clustered predominately around epithelial-associated genes/pathways (Figure 1). While our small sample size does not cover the heterogeneity and spectrum of asthma severity, our data is consistent with studies of other epigenomic mechanisms (10, 13, 14) and adds to growing evidence that chronic inflammatory airway disease is characterized by distinct changes to the epigenome.

By profiling an enhancer-associated histone modification we could identify for the first time, to our knowledge, airway epithelial cell- and asthma-associated SEs (Figure 2). While validation in a larger cohort is needed, we found that asthma-SEs encompassed transcription factors (*TP63*)(41), non-coding RNAs regulating cell function following viral infection (e.g. *miR-31*)(48) and enzymes important in lipid metabolism (e.g. *PTGS1, NOX4*, Table S4)(49). Given lipid mediators are key components of the inflammatory response and targets of current (e.g. montelukast) and novel asthma therapies (e.g. fevipiprant)(50), further investigations are required to determine the consequence of epigenome perturbations surrounding enzymes responsible for mediator synthesis.

We sought to identify the molecular components associated with changes in H3K27ac in asthma BECs and augmented TF motif enrichment with pathway analyses and protein expression data (Figure 3). Using this approach, we identified a transcription factor signature that was enriched for factors known to drive airway epithelium lineage determination and pathology (e.g. FOX- and P53-family members) and inflammation (AP-1). We postulate that the TFs identified in our analyses reflect the ‘master TFs’ of discreet cell populations comprising the airway epithelium and how this admixture may differ in people with asthma. Supporting these speculations, we found a number of TFs were encompassed by BEC-SEs including TP63, a well-defined marker of basal cells(51) and TP73, a TF with a key role in mucociliary development (44)(Figure S2C) and observed enrichment for FOXI1 sites in asthma DERs, a TF shown to characterize the recently identified ionocytes (52, 53).

While direct relationships have yet to be investigated, we believe that binding sites of the TFs identified represent an additional set of regulatory domains that cluster together to influence the expression of genes involved in key processes of the airway epithelium such as cilia formation (e.g. TP73 and FOX/RFX-family member interactions(54)), cell integrity and cell cycle progression(Figure 3F). However, we speculate that in asthma these same TFs may facilitate disease-associated alterations to the epigenome by themselves possessing pioneer or chromatin remodeling capacity (e.g. FOX-A members)(55), exhibiting differential H3K27ac at their respective gene loci (e.g. loss of H3K27ac across TP73, Figure S2C) and having altered gene expression profiles in asthmatic epithelial cells(34). These speculations would also suggest that some of the characteristics of the asthmatic airway (e.g. defects in differentiation/repair) involve an epigenomic component and could explain why these characteristics are maintained *ex vivo* (41). Consequently, genome-wide profiling studies are therefore needed to establish whether the TFs enriched in Asthma DERs represent the drivers initiating and exploiting changes we observed in the histone landscape of asthma BECs.

We employed a stringent computational method to analyze our H3K27ac ChIP-Seq data to identify regions which were differentially acetylated in asthma BECs and integrated these data with those derived from the largest studies to date of DNA methylation and gene expression in the airway epithelium of people with asthma. We found enrichment and a weak but statistically significant correlation between H3K27ac and asthma-associated CpG methylation in bronchial brushings by Nicodemus-Johnson *et. al*(13)(Figure 4B). However, this relationship accounted for less than 10% and 5% of differentially methylated CpGs and asthma DERs respectively, likely reflecting the CpGs assayed on the 450K DNA methylation array occur predominately in TSS-associated CGI’s rather than intergenic regions where most of the asthma-associated changes in H3K27ac occurred (Figure 1C). Similarly, we found that the gain and loss of H3K27ac was associated with concurrent changes in asthma-associated gene expression reported in bronchial brushings by Modena *et. al* (34). However, while traditionally histone acetylation is a hallmark of transcriptional activation, we found discordance in this association across many genes (Figure 5C).

While we identified some relationships from our data integration approach, we relied on data generated from different assays (ChIP-Seq Vs Microarray) and derived from different studies, each containing their own heterogeneous healthy and asthmatic populations. Additionally, the challenge of integrating transcriptome with epigenetic datasets is that the transcriptome can be quickly influenced by exposure to stimuli, producing different quantities of gene transcripts across a population of cells (or donors) depending on location, timing and exposure intensity. In these situations, the epigenomes of the cells sampled may be altered in a disease-specific manner but will not correlate with gene expression profiles. Furthermore, control of the transcriptome is extremely complex with each transcript potentially having multiple distal and proximal regulatory domains. Equally, the epigenome is under its own complex control, with the removal and deposition of epigenetic marks being tightly regulated (e.g. the DNA methylome may have multiple CpGs in any locus which may be differentially methylated). Further confounding these relationships is the influence of underlying genetics and the presence or absence of disease-associated polymorphisms.

However, it is becoming apparent that the complexities between epigenetics, genetics and transcriptome may be clarified by taking higher order chromatin architecture into account(56). Indeed work by Schmiedel *et. al*(46) reported that asthma-risk alleles perturb long-range interactions involving *ORMDL3* in CD4^+^ cells. Likewise, DNA methylation at asthma-associated SNPs are also implicated with long-range interactions across 17q21 locus (13). When we integrated 3D architecture data into our own analyses, we observed long-range interactions connected SNPs, DERs and BEC-SEs across the 17q21 (Figure 4G) and CLCA1 loci (Figure 5E).

Our observations add to growing evidence that in order to clarify the complex molecular relationships that underpin asthma, future studies should seek to firstly address disease heterogeneity by generating matched genome-wide datasets across all study volunteers and subsequently integrate genotype, epigenomic, transcriptomic and 3D architecture data and accurately determine asthma-associated quantitative trait loci (56).

To validate our ChIP-Seq analysis and identify a functional consequence of H3K27ac in asthma, we employed a CRISPR-Cas9-based approach to recapitulate the epigenome of asthma BECs in human embryonic kidney (HEK293T) cells *in vitro*. We targeted acetylation to asthma DERs at *TLR3* and found that gene expression could be induced in an acetylation and DER-specific gRNA manner, providing evidence that asthma-associated transcriptional profiles are driven in part by aberrant acetylation (Figure 5G).

We attempted to repeat these experiments in more relevant airway cells lines (e.g. BEAS-2B, 16HBE) but could not induce similar changes in gene expression (data not shown). Studies by Klann *et. al* (57) have indicated the ability to manipulate the epigenome using dCas9 approaches can be cell-type specific, suggesting that when a target region in a population of cells is already acetylated, dCas9-based epigenome editing may only produce a modest effect due to lack of substrate availability. Furthermore, the combination of other regulatory domains/factors may be needed to define expression capability of particular transcripts. Indeed, we found targeting an asthma-SEs did not result in induction of gene expression in HEK293T cells (Figure S2G). These data suggest that some enhancers are cell-specific or require additional stimuli to be functional (e.g. altering 3D architecture bringing enhancers/TF complexes to gene promoters).

Application of dCas9-based epigenome editing to add/remove epigenomic marks (e.g. histone modifications, DNA methylation) in a site-specific manner will be most effective if target nucleosomes are devoid of any such modification. Unfortunately, recapitulation of the entire epigenetic landscape of target genes is impossible at present, partly because the complete repertoire epigenomic marks for any gene is not understood and partly because it is currently not possible to differentially target individual nucleosomes or individual CpG dinucleotides. Regardless, dCas9-based epigenome editing provides a tremendous tool to clarify the functional consequences of asthma-associated changes in the epigenome(58).

It remains to be determined what the trigger(s) mediating the changes in the epigenomic landscape we identified in asthma is/are (Fig 1). Our asthmatic volunteers had not had exacerbation for at least 6 weeks prior to sampling so any differences observed would represent an established ‘memory’ of disease that is not diminished by *ex vivo* culture. We feel that epigenome reorganization may have developed over time and question where, when, and under what circumstances does epigenome reorganization originate? Future longitudinal studies involving larger samples numbers and covering the spectrum of asthma severities are needed to determine when the epigenome is altered in asthma and if there are key ‘epigenomic windows’ of disease development that might benefit through therapeutic intervention.

## Methods

### Patient Recruitment

Volunteers with and without asthma were recruited as previously described(23). None were smokers or had had asthma exacerbations or any respiratory tract infections in the preceding 6 weeks (Table S1). Asthma was defined as a physician’s diagnosis of asthma, and volunteers with asthma had airway hyper-responsiveness (provocative concentration (PC_20_) of histamine required to reduce FEV_1_ by ≥20% of ≤8 µg/mL, an Asthma Control Questionnaire(24) score >0.75 and were on treatment with inhaled corticosteroids (ICSs) or a combination inhaler (long-acting β agonist ⍰ + ⍰ ICS). Healthy controls had a PC_20_ histamine >8 µg/mL. All volunteers with asthma were atopic and all healthy controls non-atopic as determined by skin prick testing (at least one positive skin prick test to a panel of 10 aeroallergens, including grass).

### Primary bronchial epithelial cell (BEC) culture

Bronchial brushings were obtained from volunteers by fibre-optic bronchoscopy using a Keymed BF260 bronchoscope (Olympus, Essex, UK) and 5 mm sheathed endobronchial brushes (Olympus BC-202D-5010) in accordance with British Thoracic Society guidelines(25). Freshly brushed BECs were removed from brushes by agitation and seeded into supplemented bronchial epithelial growth medium (BEBM, Lonza) in a T25 flask. Cell culture was performed as described previously(5), and seeded at passage 2 onto 10mm culture dishes (Primaria, Corning) and cultured until 80% confluent.

### Chromatin immunoprecipitation

BECs were trypsinized, lysed in hypotonic/mild detergent buffer (15Mm Tris-HCL pH7.5, 60mM KCL, 15mM NaCL, 5mM MgCl_2_ 0.1mM EGTA, 5mM DTT, 0.4% IGEPAL-CA) and nuclei isolated through sucrose density gradient (0.3/1.2M sucrose, 20,000g, 20min, 4°C)). Viability of nuclei was assessed via trypan blue staining. Nuclei were treated with micrococcal nuclease (10U, NEB) for 10 minutes at 37°C, re-pelleted and supernatant containing mononucleosomes stored at 4°C. Mononucleosomal fractions were incubated with 4μg of anti-H3K27ac antibody (Abcam, Ab4729) and 25μL Protein G dynabeads (Life Technologies) in modified RIPA buffer over night at 4°C. Bound complexes were washed and eluted and ChIP DNA extracted using phenol:chloroform/ethanol precipitation.

### Sequencing and mapping

Libraries were prepared using half total volume of eluted ChIP DNA and NEBNext® DNA Library Prep Master Mix Set and Multiplex Oligos for Illumina® (New England Biolabs). Library quality was assessed using Bioanalyzer 2100 High Sensitivity DNA Gels (Agilent). Libraries were subject to 50bp single end read sequencing on the HighSeq 2500 (Illumina) in rapid run format and reads were aligned to Human genome hg19 using Bowtie2 (Galaxy v2.2.6(26), Table S2). ENCODE and BEC-associated blacklist regions (including chr*N*, chr*Un* and chrM) were subtracted from each BAM file prior to downstream analysis using the *intersect -v* function in bedtools (Galaxy v2.1.0). For visualization, Healthy (n=3) and Asthma (n=4) BAM files were merged using samtools (Galaxy), Input BAMs subtracted (-*bamCompare*) and BigWig’s generated and normalized to reads per kilobase per million mapped reads (RPKM, -*bamCoverage*, deepTools) (27).

### Differential Analysis and super-enhancer identification

Peaks were called for merged Healthy, Asthma and individual volunteer BAM files using MACS2 (p<5e-4, Galaxy v2.1, Table S2). Differentially enriched regions (DERs) between Asthma (n=4) and Healthy BECs (n=3) were identified from a consensus set of peaks (n=42,168) using DiffBind (28)(*Summits =500, FDR Threshold=0.05*, Table S3). Significance of DERs genome-wide was visualized with Manhattan plots generated using qqman and genomic distribution of DERs determined using CEAS (CISTROME v1.0.0). DERs were annotated using *ROSE_geneMapper.py* (see below) and HOMER(29).

Typical and Super enhancers (SEs) as defined by Hnisz et al.(16) were identified and annotated using Rank Ordering of Super Enhancers algorithm (*ROSE_main.py, -t 2000 and ROSE_geneMapper.py, searchwindow=default* respectively, Table S4)(16). Individual volunteer SEs were compared using *merge -o distinct, count* function in bedtools to identify SE categories. Briefly, to strike a balance between small volunteer numbers and disease heterogeneity, Common SEs were defined as those being present in n=4/7 volunteers, Healthy SEs; n=2/3 healthy volunteers and Asthma SEs; n=3/4 asthma volunteers and no further overlap with any other SE category. dbSUPER(17) overlap analysis (default settings) was used to determine similarity of BEC-SEs with other cell type-SEs and heatmaps were produced using Morpheus (http://software.broadinstitute.org). BEC-SEs were intersected with Asthma DERs and coverage of DERs across BEC-SEs determined using the *intersect* and *coverageBed* options in bedtools. BEC-SE ‘metagene’ H3K27ac plots were generated using *plotProfile* in deepTools.

### Transcription factor motif enrichment

Motif enrichment in DERs (n=4,321) and peaks within BEC-SEs (n=12,457) was conducted using HOMER (*findMotifsGenome.pl -mknown=1HOCOMOCOv11 core HUMAN mono homer format 0.001.motif* and options for DERs (*-size given, -h -bg=all test peaks*) and BEC-SEs (*-size 500*)(29, 30). We then selected enriched TFs (P<0.05) associated with epithelial processes via Ingenuity® (see below) and/or had motifs enriched in BEC-SEs and/or had protein expressed in the respiratory epithelium for further analysis (n=201, Figure S2A-B). Protein expression and tissue specificity of TFs was determined via Human Protein Atlas (normal tissue data v.16.1)(31). Heatmaps were produced using Morpheus. Published TF ChIP-Seq datasets were download from ChIP-Atlas (32) and merged to create a consensus list of bona-fide TF binding sites and intersected with DERs using bedtools. HOMER (*annotatePeaks.pl - m=HOCOMOCOv11 core HUMAN mono homer format 0.001.motif*) was used to predict binding sites for those TFs with no ChIP-Seq data available. Binding site enrichment was determined using Fishers Exact test and plotting odds ratio (OR, +/-95% confidence intervals) with the consensus peak set identified in DiffBind analysis (n=42,168 peaks) used as background. H3K37ac enrichment profiles ±0.55kb around intersecting DER center were plotted using *plotProfile* in deepTools.

### Pathway analysis

All pathway and biological process enrichment analyses and SE-associated gene categorizations were conducted using Ingenuity® Pathway Analysis Software (QIAGEN).

### Asthma DNA methylation and SNPs

DER overlap with ENCODE CpG islands was conducted with bedtools and enrichment determined using Fishers Exact test as above. H3K37ac enrichment profiles ±0.5kb around intersecting DER center were plotted using deepTools. Percentage of CpG content for asthma DERs was determined using HOMER and correlation analysis with differentially methylated CpGs identified in asthma BECs by Nicodemus-Johnson *et al*.(13) conducted using GraphPad Prism v.8.1. DERs were intersected with Asthma SNPs compiled by Seumois *et al*.(10). Relationship between SNPs and distance to DERs/enhancers (i.e. peaks >2kb from TSS) was determined using *closestbed -d ref* function in bedtools and binning distances using the *frequency* option in GraphPad Prism v.8.1.

### 3D chromosomal interactions

To investigate relationships between long-range interactions, asthma SNPs and DERs, chromosomal interaction data (Hi-C) was accessed via 3D Interaction Viewer and Database(33). Since no chromosomal interaction data is available for normal airway epithelial cells, we accessed datasets from whole lung tissue. Search parameters were; *Bait=ORMDL3/CLCA1* TSS, *Interaction range=500kb*, *TAD=TopDom (w=20)*. Interactions were intersected with asthma SNPs/DERs using bedtools and distance-normalized interaction frequencies of overlapping 5kb bins used to determine strength of SNPs/DER-containing interactions.

### Transcriptomics

We investigated gene expression using published microarray dataset GSE63142(34), the largest study to date of bronchial brushings from healthy controls and mild/moderate and severe asthmatics. Since our ChIP-Seq dataset was derived from mild asthmatics, we focused on mild/moderate asthmatics (n=72) and controls (n=27) and undertook differential gene expression analysis using Partek Genomics Suite (v6.4). After correction for batch effects, ANOVA analysis identified n=4,669 differentially expressed (DE) genes (6,080 probes, P<0.05). Relationships between differential gene expression and asthma DERs were determined using BETA (Cistrome v1.0.0, *geneID=Refseq, genome=hg19, peaks=30,000, TSS distance=50,000, CTCF=True, significance=0.05*)(35). Briefly, by assigning a rank number based on the distance between asthma DERs and DE genes (x axis) and plotting them versus the proportion of genes with ranks at or better than the x axis value (y axis), BETA identifies the potential activating/repressing function of asthma DERs on gene expression.

### Targeting acetylation to asthma DERs

Plasmids encoding active and mutant forms of a deactivated-Cas9-histone acetyltransferase P300 fusion protein (pcDNA-dCas9-p300 Core, pcDNA-dCas9-p300 Core-D1399Y) and guide RNA only expression vector (phU6-gRNA) were gifts from Charles Gersbach(22)(Addgene #61357, #61358 and #53188 respectively). DER-specific gRNAs were designed by submitting DER coordinates or sequence to the CRISPOR (http://crispor.tefor.net) and Breaking-Cas(36) servers respectively and selected based on overlap with BEC DNAse hypersensitivity sites (ENCODE). Control gRNAs to IL1RN were described previously(37). gRNAs were annealed and cloned into phU6-gRNA via *BbsI* restriction sites (NEB #R0539S). All plasmids were transformed into OneShot®TOP10 competent cells (ThermoScientific), cultured overnight and extracted using Endo Free® Plasmid Maxi or QIAprep® Spin Miniprep (gRNAs) kits (QIAGEN). gRNAs sequence and locations are listed in Table S5.

### Cell culture and transfections

HEK-293T cells were maintained in DMEM media supplemented with 10% fetal calf serum and penicillin/streptomycin at 37°C 5% CO_2_. 2×10^5^ cells were reverse transfected with 1μg of dCas9-P300 constructs and 125ng of equimolar pooled or individual gRNAs using Lipofectamine 3000 according to manufacturer’s instructions (Thermoscientific). Cells were harvested at 48hrs and RNA extracted (RNAeasy® Mini Kit, QIAGEN). Gene expression was determined via Taqman qPCR assays (Table S5) and run on the Applied Biosystems ViiA^TM^ 7 Real Time PCR system (ThermoFisher Scientific).

### Glucocorticoid receptor ChIP-Seq

FASTQ files from dataset GSE79803(38) was downloaded from SRA (SRP072707) and processed as outlined above. DERs were intersected with dexamethasone (DEX)-responsive GR sites as described by Kadiyala *et al*.(38) using bedtools and GR enrichment profiles ±5kb around intersecting sites plotted with deepTools.

## List of abbreviations

3D: three dimensions
BECs: bronchial epithelial cells CGI’s – CpG Islands
ChIP-Seq: Chromatin-immunoprecipitation coupled with high-through-put sequencing.
CRISPR-Cas9: Clustered, regularly interspaced, short palindromic repeats.
DERs: differentially enriched regions
DEX: dexamethasone
ENCODE: Encyclopedia of DNA Elements consortia
gRNA: guide RNA
H3K27ac: Histone H3 Lys27 acetylation
H3K4me2: Histone H3 Lys4 dimethylation
kb: kilobase
RPKM: reads per kilobase per million mapped reads
SEs: super enhancers
SNPs: single nucleotide polymorphisms
T2: Type 2 airway inflammation
TFs: transcription factors
TSSs: transcriptional start sites

## Declarations

### Ethics approval and consent to participate

This study was approved by the London Bridge Research Ethics Committee (reference 10/LO/1278) and was carried out in accordance with the Declaration of Helsinki and Good Clinical Practice guidelines. Informed consent was obtained from all subjects prior to their participation.

### Consent for publication

### Availability of data and material

All data has been deposited on gene expression omnibus (GSE109894).

### Author Contributions

J.D. performed the clinical aspects of the study. S.L.J. supervised clinical aspects of the study.

M.R.E performed clinical sample processing and culture. P.M., A.K, J.W. performed ChIP Experiments.

P.M. and P.L. analyzed ChIP-Seq data. P.M., M.K. performed dCas9 CRISPR work.

P.L., D.J.C., P.M, J.S., R.S., A.V.O., M.R.E and S.L.J conceived and designed the study.

## Acknowledgements

The authors wish to thank Anthony Gerber for glucocorticoid receptor Chip-Seq data and staff within the BRC Genomics platform at the Biomedical Research Centre of Guy’s and St Thomas’ NHS Foundation Trust for assistance with sequencing.

## Supplementary Figure legends

**Figure S1:**
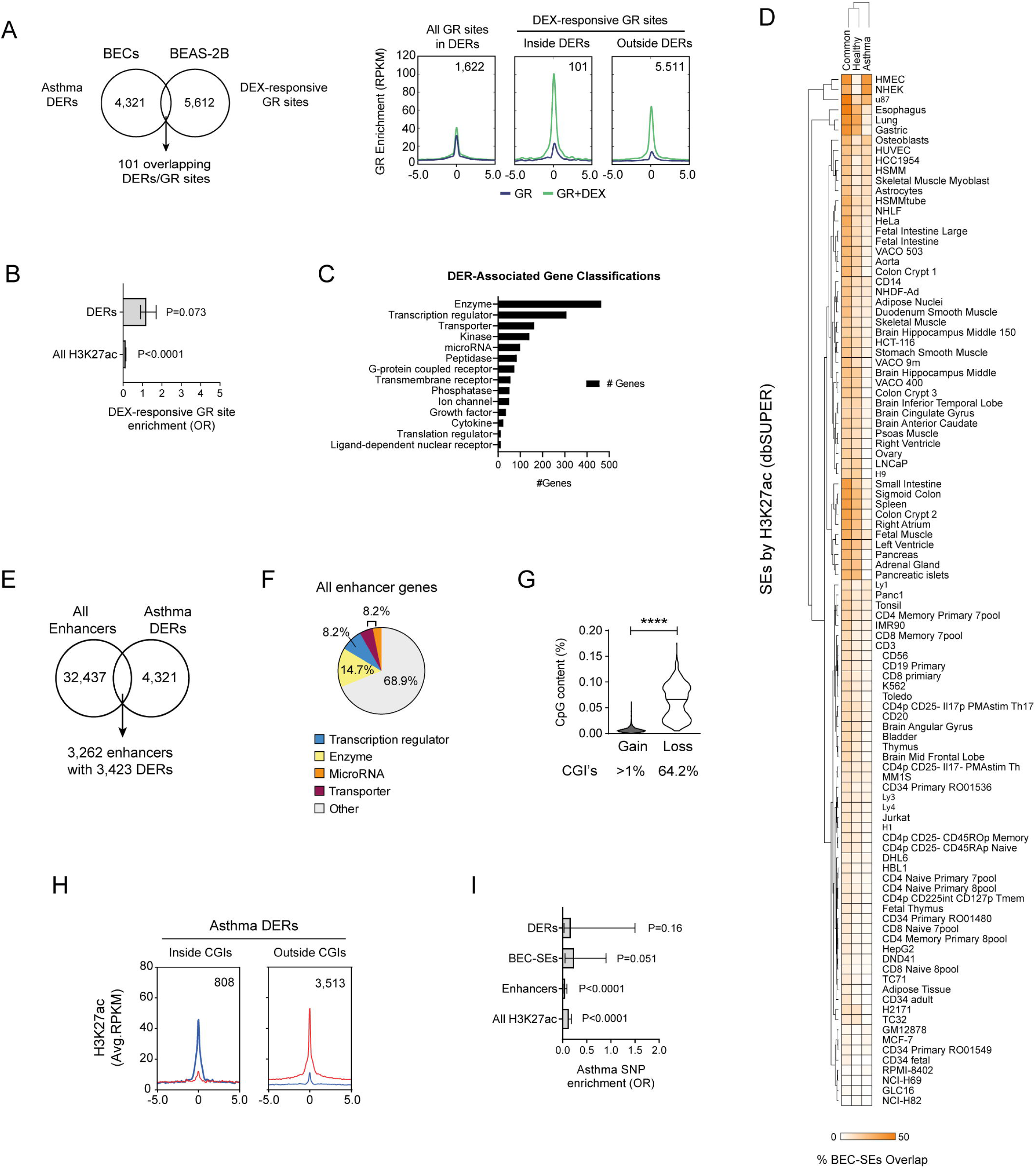
(A) Overlap between asthma DERs and dexamethasone (DEX)-responsive glucocorticoid receptor (GR) binding sites identified in the BEAS-2B airway epithelial cell line. Profile plots depict GR enrichment ±5kb around GR binding sites overlapping (inside) and non-overlapping DERs (outside). (B) Enrichment of DERs at DEX-responsive GR sites (Fishers odds ratio +/-95% CI). (C) Ingenuity classification of DER-associated genes. (D) Relationship between SEs identified in airway epithelial cells compared to SEs identified via H3K27ac ChIP-Seq in other cell types(17).(17). BEC-SEs were most closely related to other mucosal/epithelial cell-type SEs (e.g. oesophagus, lung, HMEC). (E) Overlap between all enhancers (typical and super) identified via MACS2/ROSE and asthma DERs identified via DiffBind. (F) Categorization of all ‘typical’ enhancer genes (i.e. non-BEC-SEs associated) identified in BECs via Ingenuity. (G) Asthma DERs exhibited differences CpG content with loss of H3K27ac in asthma more likely to overlap CpG Islands (CGI’s, mean ±max-min, **** *P*<0.0001, Wilcoxon test on ranks). (H) Profile plot depicting H3K27ac enrichment of DERs in relation to CpG islands (CGIs) across Healthy (blue) and Asthma BECs (red). (I) Enrichment of asthma SNPs across DERs and other features of H3K27ac in BECs (Fishers odds ratio +/-95% CI).

**Figure S2:**
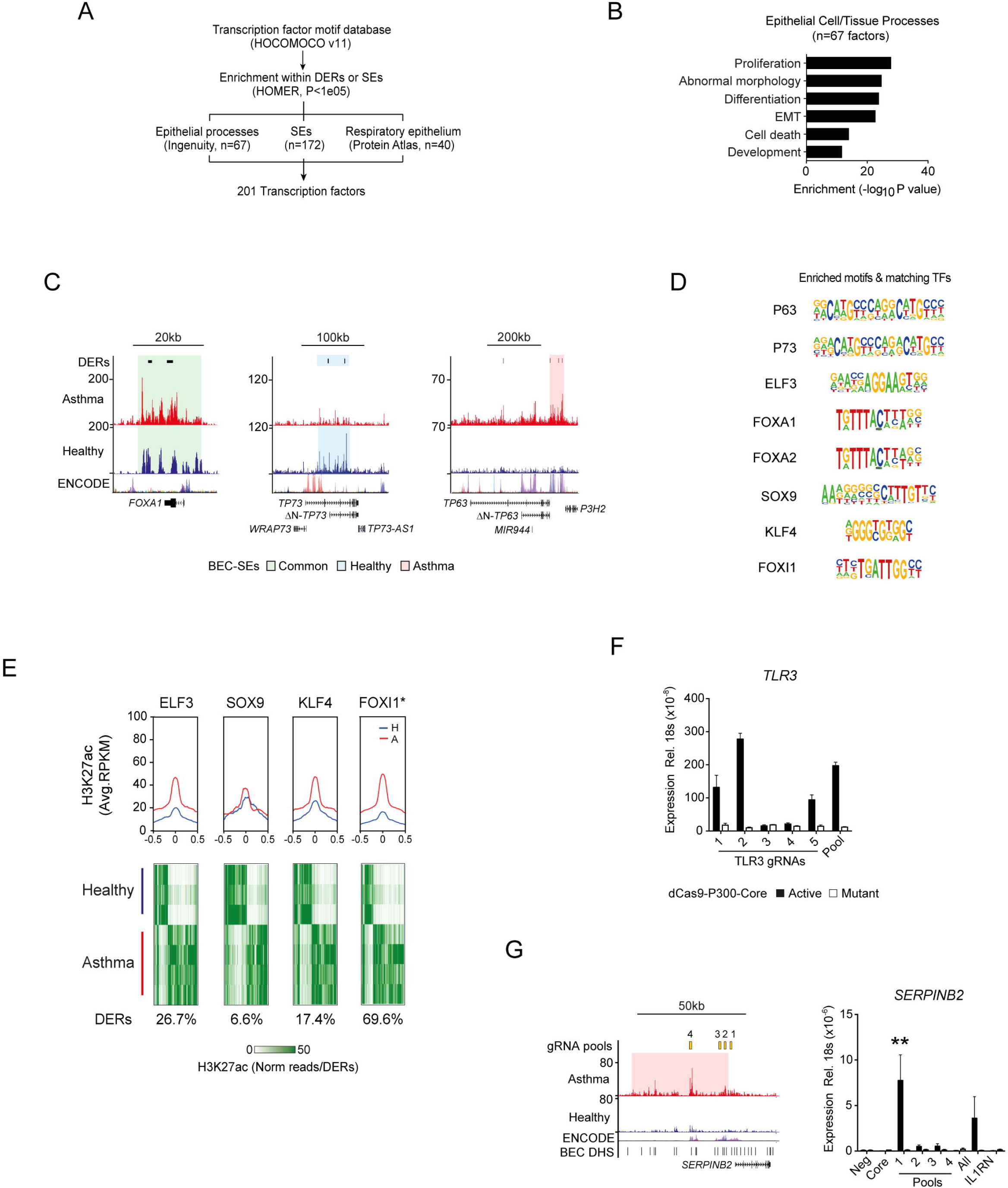
(A) Work flow used to refine motif enrichment analysis and focus on enriched transcription factors (TFs) important in airway epithelial biology and BEC-SEs. (B) Ingenuity analysis indicating TFs with motif enrichment in asthma DERs and BEC-SEs were associated with various processes in epithelial biology including epithelial to mesenchymal transition (EMT). (C) Genome tracks depicting H3K27ac across loci encoding epithelium-dominant TFs encompassed by BEC-SEs and asthma DERs. Asthma=red, healthy=blue. (D) Enriched motifs and matching TFs identified by HOMER (see methods). (E) H3K27ac profiles of healthy (blue) and asthma (red) BECs at epithelium-dominant TF binding sites identified via ChIP-Seq and those predicated by HOMER (*; see also Figure 3D) (F) *TLR3* gene expression could be induced with single DER-specific gRNAs in an acetylation-dependent manner (n=3, Mean ±SEM). (G) Genome tracks depicting H3K27ac in BECs across the *SERPINB2*. Acetylation (dCas9-P300-Core) was targeted to the TSS (pool 1) of *SERPINB2* and upstream within an asthma-associated SE (pools 2-4) using gRNAs (yellow boxes top tracks). No induction of gene expression was observed when targeting distal enhancer elements. Guide RNAs to *IL1RN* used as controls. Core - only P300 constructs transfected (n=3, Mean ±SEM, ** *P*<0.01, ANOVA/Tukeys multiple comparisons).

**Table S1:** Clinical characteristics of study volunteers. ACQ – asthma control questionnaire, FEV_1_ – forced expiratory volume in one second, PC_20_ - provocative concentration of histamine to cause a 20% decrease in FEV_1_. SPT – skin prick test.

**Table S2:** Statistics for H3K27ac ChIP-Seq data and analysis.

**Table S3:** Differentially enriched regions (DERs) of H3K27ac in Asthma BECs. Table lists all DERs FDR *P*<0.05, associated genes and H3K27ac enrichment across all study volunteers. Additional annotations by HOMER are included.

**Table S4:** Super enhancers of the airway epithelium.

**Table S5:** Sequence and location of DER-specific and control guide RNAs and Taqman assays used in this study.

